# Conservation and host-specific expression of non-tandemly repeated heterogenous ribosome RNA gene in arbuscular mycorrhizal fungi

**DOI:** 10.1101/2020.05.14.095489

**Authors:** Taro Maeda, Yuuki Kobayashi, Tomomi Nakagawa, Tatsuhiro Ezawa, Katsushi Yamaguchi, Takahiro Bino, Yuki Nishimoto, Shuji Shigenobu, Masayoshi Kawaguchi

**Affiliations:** Division of Symbiotic Systems, National Institute for Basic Biology, Nishigonaka 38, Myodaiji, Okazaki, Aichi, Japan; Graduate School of Agriculture, Hokkaido University, Sapporo, Japan; Functional Genomics Facility, National Institute for Basic Biology, Nishigonaka 38, Myodaiji, Okazaki, Aichi, Japan

## Abstract

The ribosomal RNA-encoding gene (rDNA) has a characteristic genomic nature: tens to thousands of copies in a genome, tandemly repeated structure, and intragenomic sequence homogeneity. These features contribute to ribosome productivity via physiological and evolutionary processes. We reported previously the exceptional absence of these features in the model arbuscular mycorrhizal (AM) fungus *Rhizophagus irregularis.* Here we examine the phylogenetic distribution of the exceptional rDNA features in the genus *Rhizophagus* via improving the genome sequence of *R. clarus.* Cross-species comparison indicated similarity of their rDNAs not only in the genomic features but also in the distribution of intragenomic polymorphic sites on the paralogs. Ribosomal RNA comprises multiple domains with different functions. The two *Rhizophagus* species commonly exhibited a variation enrichment site, ES27L, which is related to translational fidelity and antibiotic sensitivity. Variation enrichment on ES27L has not been observed in other organisms lacking the three rDNA features such as malaria parasites and *Cyanidioschyzon merolae.* Expression profiling of rDNAs in *R. irregularis* revealed that rDNA paralogs are expressed differently in association with host plant species. Our results suggest a broad distribution of the disarranged rDNA across AM fungi and its involvement in the successful association with the broad range of host species.

## Introduction

Ribosome heterogeneity was first proposed by Francis Crick as the “one gene–one ribosome-one protein” hypothesis in 1958 [1]. This hypothesis, where a different type of ribosome translates each protein, vacillated within a few years, and the idea that “all ribosomes are exactly the same” became prevalent [2]. However, recent evidence of heterogeneous ribosomes in various organisms (e.g., humans, mouse, yeast, *Arabidopsis,* fruit flies, zebrafish, and malaria parasites) rekindled the notion that ribosome is an additional regulatory layer of gene expression [2–4]. Although a number of studies have reported the correlation between tissue- and cell stage-specific phenotypes and specialized ribosomes, the detailed expression control mechanisms and evolutionary contributions of this heterogeneity remain ambiguous [2–4]. Notably, the importance and distribution of the heterogeneity of the ribosomal RNA (rRNA) remain an open question. The reliable sequencing and mutant/transformant construction of the 18S–5.8S–28S ribosomal RNA-encoding loci (48S rDNA) have been technically challenging due to their large copy number (CN) and tandem repetitive structure (TRS) [2].

In the previous genomic study of the 48S rDNA in the model arbuscular mycorrhizal (AM) fungus *Rhizophagus irregularis* DAOM-181602, we unexpectedly found (1) a small copy number (around ten copies), (2) the absence of TRS, (3) and intragenomic heterogeneity [5]. AM fungi belong to the subphylum Glomeromycotina [6] and form symbiotic associations with most land plants [7, 8], and to date, up to 300 species have been described [9]. The association was established in the early Devonian and contributed to plant terrestrialization via enhancing nutrient uptake [10, 11]. AM fungi colonize plant roots and construct extensive hyphal networks in the soil and deliver essential nutrients to the host plant. AM fungi show no apparent host specificity; they are capable of colonizing different plant functional groups, that is, autotrophic and heterotrophic plants, and connect them via the underground mycelial networks in the field [12, 13].

According to the recent ideas for the physiological contribution of ribosome heterogeneity in eukaryotes [2–4], it is hypothesized that the ribosome polymorphisms in *R. irregularis* assist adaptation to diverse environments and facilitate a broad host range via translational modifications [5]. The absence of the TRS of rDNA may allow us to elucidate not only the evolutionary mechanisms underlying the conservation of the TRS in the majority of eukaryotes but also those for overcoming the difficulty in rDNA mutation/transformation construction. However, the unique features of AM fungal rDNA have only been discovered very recently [5, 14], and the extent of the features in the Glomeromycotina still remains unexplored. Although genome sequences have recently been published in several species, including *R. clarus, Glomus cerebriforme, R. diaphanous, Gigaspora rosea, G. margarita,* and *Claroideoglomus claroideum* [15–18], the genomic structure of rDNA has been unclear except in *R. irregularis*, which is due to, at least partially, the difficulty in constructing genome assemblies long enough for the analysis of the rDNA structure.

To examine the nature of AM fungal rDNA in greater detail, we improved the genome data of *R. clarus* HR1 and analyzed whether *R. clarus* HR1 shares the three rDNA features and polymorphic sites with *R. irregularis* via cross-species comparison. Subsequently, we performed PacBio circular consensus sequencing (CCS) [19], which generates accurate long-read sequences, for profiling the expression of different rDNA paralogs in association with the environmental condition.

## Results

### Improved *R. clarus* genome assembly indicates high similarity in rDNA genomic structure between *R. irregularis*

Here, we obtained an improved *R. clarus* genome assembly covering all of the 48S rDNA loci that were expected based on the read depth of coverage (**Fig. 1)**. The improved *R. clarus* rDNA resembled that of *R. irregularis:* it exhibited 11 copies, a non-tandemly repeated structure, and heterogeneity among the copies (**Fig. 1b**).

**Fig. 1.**
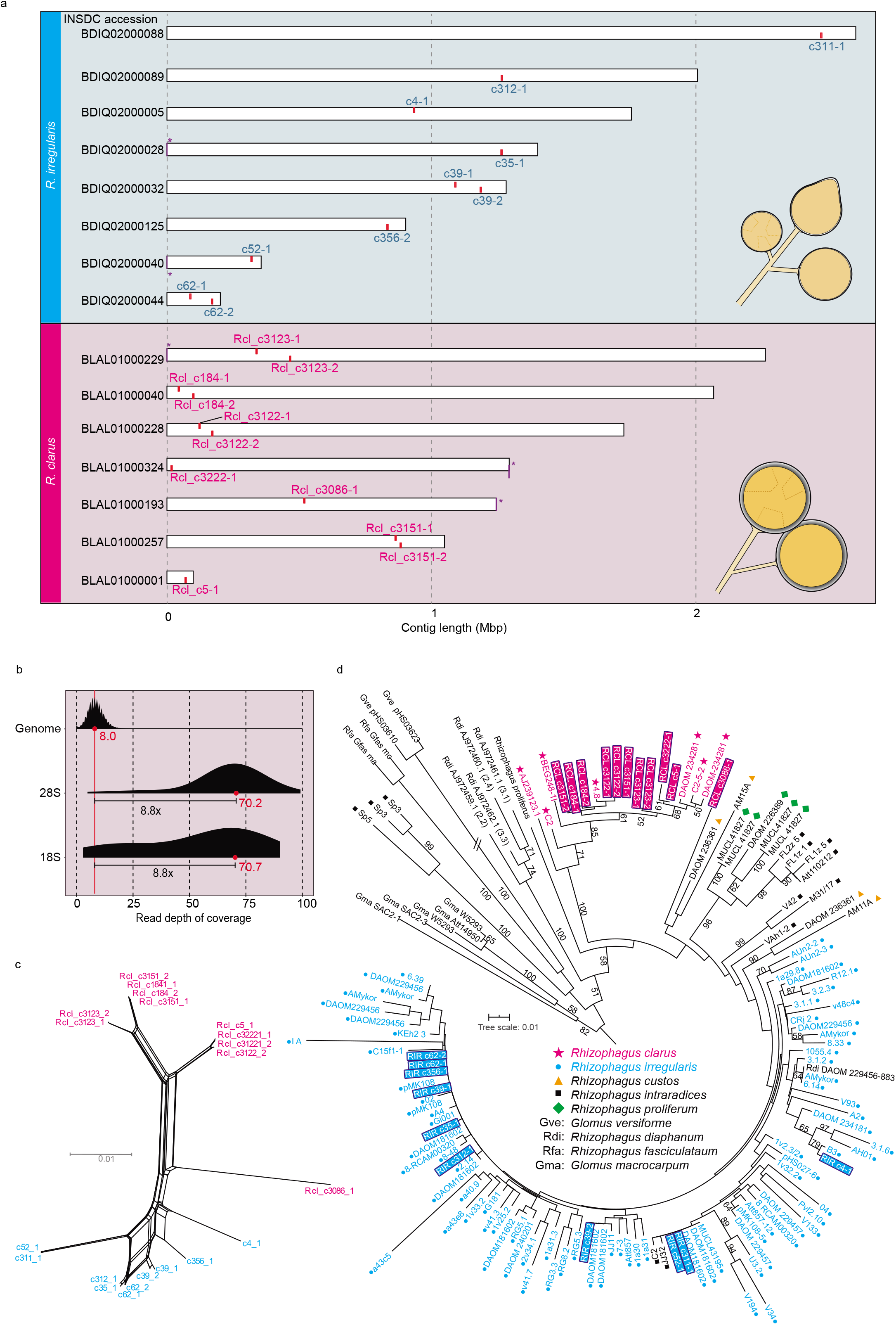
Ribosomal DNAs of *R. clarus* and *R. irregularis.* **a** Genomic positions of the 48S rDNA clusters on the constructed contigs. Rir, *Rhizophagus irregularis* (cyan); Rcl, *Rhizophagus clarus* (magenta). Red and purple bar (with asterisk) indicate 48S rDNA, and telomeric region, respectively. Contigs with no 48S rDNA were omitted from the figure. **b** Read depth of coverage of the *R. clarus* rDNA regions and whole-genomic region. Red dots indicate mode value. **c** Neighbor-net tree based on the whole sequence of 48S rDNA sequences. **d** Maximum likelihood (ML) tree based on the ITS region of the rDNAs (420 positions) (raw data, https://doi.org/10.6084/m9.figshare.11880834, 10.6084/m9.figshare.12251768, 10.6084/m9.figshare.11880780)

The total size of the *R. clarus* contigs was comparable with the estimated genome size, and the gene models covered almost all of the eukaryotic conserved gene sets. A total of 5,819,346 PacBio reads were generated, with an average length of 3.4□kb (**Supplementary Tables 1 and 2**). The assembling and the error correction resulted in a sequence dataset containing 147 Mbp (360 contigs, N50 = 1.30 Mbp, 31,233 genes) (**INSDC; BLAL01000001-BLAL01000360**). A BUSCO analysis revealed high coverage of the eukaryotic conserved gene set (95.1%, DB; eukaryota_odb9, **Supplementary Table 2**). *R. clarus* contained a telomeric region on the edge of multiple contigs as *R. irregularis* (7 in *R. irregularis* and 63 in *R. clarus).* The nucleotide sequence of telomeres was “TTAGGG,” similar to that of the majority of fungi [20]. Although the nuclear phase remains unclear in AM fungi [21], our data informed us that the AM fungus has the usual telomeric region and suggested the minimum expected number of chromosomes.

Our *R. clarus* assembly shared the three exceptional rDNA features with *R. irregularis.* RNAmmer found 11 copies of the complete 45S rDNA cluster, which was composed of 18S rRNA, intergenic spacer region 1 (ITS1), 5.8S rRNA, ITS2, and 28S rDNAs (**Fig. 1a, Supplementary Data 1, Supplementary Figure 1**). None of the *R. clarus* rDNAs formed the TRS, comprising multiple tandemly repeated units of the 45S rDNAs. Most of the 45S clusters were located on different contigs; a single copy of rDNA was detected in three contigs, and two copies were found in four contigs (Fig. 1a). In cases in which two rDNA clusters were found, the two copies resided apart from each other and did not form a tandem repeat. The internal regions contained protein-coding genes, respectively, and the two clusters were located on reverse strands from each other. Because all rDNA copies were located >10 kb away from the edge of each contig (**Fig. 1a**), the absence of TRS was unlikely to be an artifact derived from an assembly problem. We found no synteny between the two *Rhizophagus* rDNAs.

To confirm that no rDNA clusters were overlooked, we estimated the rDNA CN based on the read depth of coverage. The mapping of the genomic Illumina reads onto the selected reference sequences indicated that the average coverage depth of the consensus rDNA was approximately nine times deeper than that of the genome, suggesting that *R. clarus* possesses around nine copies of rDNAs in its genome (**Fig. 1b, Supplementary Data 2**). Thus, we considered that our genome assembly (containing 11 rDNA copies) covered almost all of the rDNA copies.

The obtained rDNAs indicated polymorphism among the 45S rDNA clusters on *R. clarus,* similar to that of *R. irregularis.* Pairwise comparisons of the 11 rDNA copies detected an average of 40.9 indels and 76.8 sequence variants, whereas one of the sequence pairs (RCL_c3122_1 and RCL_c3122_2) had identical sequences (**Supplementary Data 3**).

To examine the phylogenetic distribution of the heterogeneous rDNAs, we constructed a phylogenetic tree with previously released *Rhizophagus* ITS sequences (**Fig 1d**). Before the phylogenetic analysis, we improved the erroneous region on the previous PacBio-based *R. irregularis* genome (GCA_002897155.1) using Illumina reads. The previous genome, which was corrected on the GATK software [5], was improved via a standard correction software for long-read-based assemblies, Pilon [22]. The new assemblies obtained differed from the previous genome at 3,660,534 positions (2.4% of the alignment, **INSDC; BDIQ02000001-BDIQ02000210**). However, the ten 48S rDNA clusters exhibited no differences compared with the previous genomes. We used these rDNA regions as reference sequences hereafter. Phylogenetic analysis revealed that the rDNA polymorphisms of our *R. clarus* genome covered most of the polymorphisms reported previously for this species. Moreover, an rDNA cluster, RCL_c3086_1, established a clade with the *Rhizophagus cactus* rDNA (**Fig 1d**).

### Distribution of non-tandemly repeated rDNAs across AM fungi

We researched the extensive conservation of the *Rhizophagus*-type rDNA using previously released fungal genomes. We selected four AM fungi and 20 non-AM fungal species from the public database and obtained successive results from two AM fungi and five non-AM species (**Supplementary Data 2**). The read depth and direct search of rDNA regions showed the conservation of the copy number reduction (CNR) and the TRS-lacking features among AM fungi. In the non-AM fungi, we obtained no evidence of the presence of the *Rhizophagus* -type rDNA.

To estimate rDNA copy number variation (CNV), we compared the abundance of Illumina reads aligned to rDNA and genomic sequences [23]. Due to the difficulty of the rDNA region assembling, we presumed that the read depth gives a more reliable estimation of the rDNA CNV than does the homology-based search of the assemblies. The estimated number of 48S rDNA copies was 1 or 10 in AM species (*G. cerebriforme* and *D. epigaea)* and 71–496 in the non-AM fungi (**Fig. 2, Supplementary Data 2**). These results suggest that the CNR is not limited to the genus *Rhizophagus* but is seemingly common among AM species. In non-AM species, we found no CN under 20 (**Fig. 2)**, which is lethal for yeast [24]. Even if we included the six species that contained a single or partial rDNA gene (see Material and Methods, **Supplementary Data 2, Supplementary Figure 2**) in the analysis, no species contained over 20 rDNA copies, with the exception of *Jimgerdemannia flammicorona (Endogonales;* estimated CN, 2), which is a plant-symbiotic species [25].

**Fig. 2.**
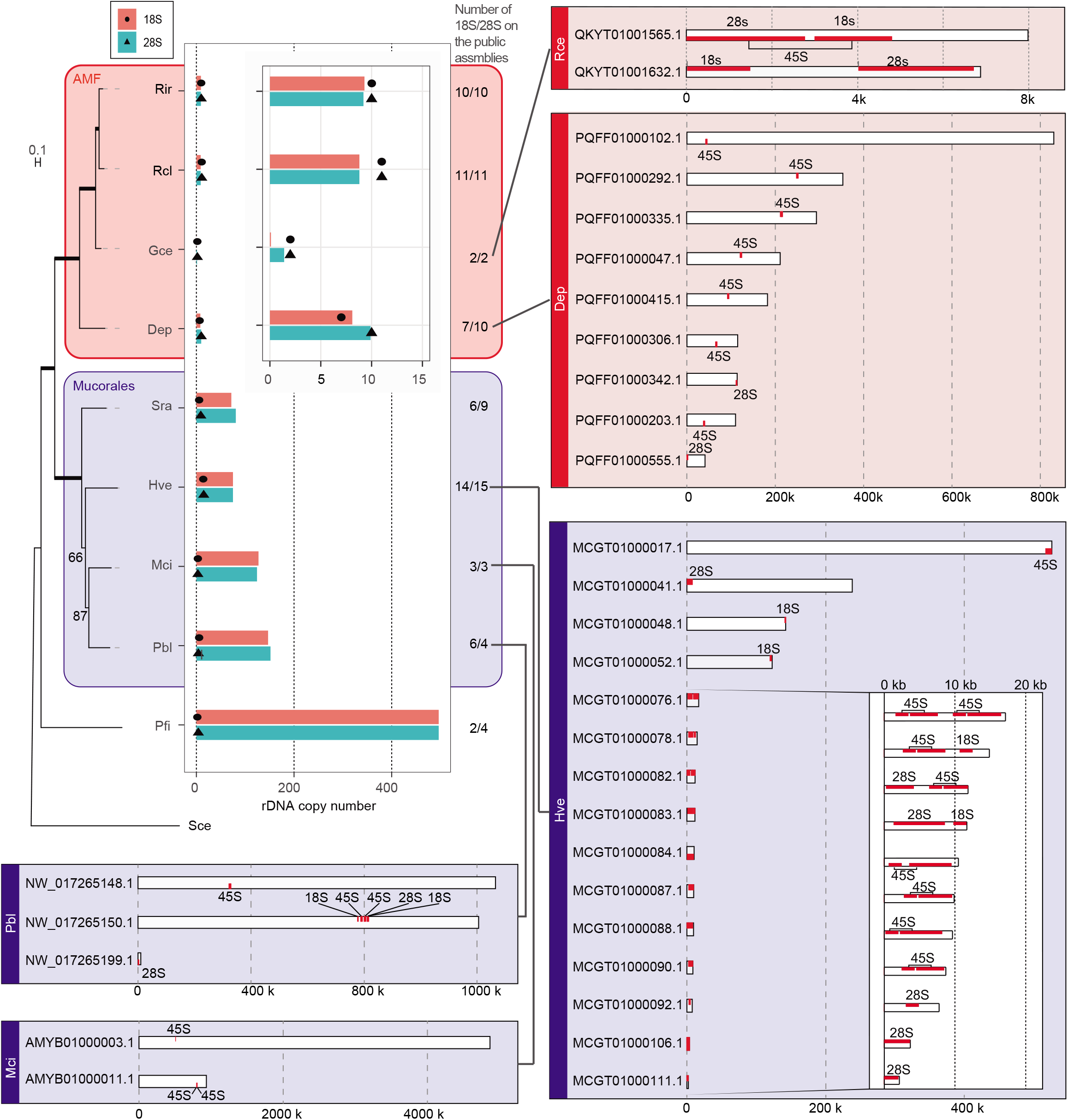
Predicted rDNA copy number and the distribution on the constructed genome data of the 18S and 28S rDNA. The colors are as in the left top box. The tree on the left was generated by the ML analysis of 96 single-copy genes. The bold line indicates the node supported with 100 bootstrap value. The species analyzed are presented using the abbreviations (Rir, *R. irregularis*; Rcl, *R. clarus*; Gce, *G. cerebriforme*; Dep, *D. epigaea*; Sra, *Syncephalastrum racemosum*; Hve, *Hesseltinella vesiculosa*; Mci, *Mucor circinelloides*; Pbl, *P. blakesleeanus*; Pfi, *Piromyces finnis*). The central bar plot shows the predicted copy number of rDNA based on the read mapping back (bar) and actually observed number in the public genome (black dot and triangle). The right plot shows the positions of the determined rDNA on each public genome sequence. Red bar indicates 48S, 18S, or 28S rDNA-encoding region. The sub-boxes indicate the expanded view of a part of the plot. Raw values of the analysis are presented in Supplementary Data 2.

Our searching of the rDNA regions on the public assemblies supported the absence of the TRS in the two AM species *(G. cerebriforme* and *D. epigaea).* The *D. epigaea* genome (GCA_003547095) had seven 48S rDNA clusters and two 28S rDNAs (**Fig. 2**). The depth-based CN, i.e., ten copies, suggested that the public genome contains the majority of the rDNA copies in *D. epigaea.* We cannot deny the possibility that the two 28S copies, which were located near the edge of contigs (1,760 bp and 522 bp), are part of the TRS structure. However, the depth-based CN suggests that even if the two 28S built the TRS on the inter-assembly region, these overlooked TRS would be short (under three copies). Another AM fungus, *G. cerebriforme* (GCA_003550305), had two clusters of 45S rDNA on the different contigs. Although these clusters were located near the edge of the contigs (1–3, 178 bp), the depth-based CN (one copy) denied the presence of a long TRS structure on the genome (**Fig. 2, Supplementary Data 2**).

In the five non-AM species, we obtained no evidence of the TRS-lacking feature. The public assemblies contained only part of the rDNA regions at 2–15 copies, and the rDNAs were located on the edge of assemblies (**Fig. 2, Supplementary Data 2**). Interestingly, two *Mucorales* species *(Phycomyces blakesleeanus* and *Mucor circinelloides)* contained one 45S rDNA located >10 kb away from the edge of the assembly (**Fig. 2**), respectively. Although the depth-based CN (*P. blakesleeanus,* 147–151 copies; *M. circinelloides*, 124–127 copies) indicated that the public genome data did not contain the majority of the rDNA regions, we found at least one TRS-lacking rDNA on this genome (**Fig. 2**). In an *Endogonaceae* species, *J. flammicorona,* the public genome had only one 28S rDNA. Although the read depth analysis indicated the presence of a *Rhizophagus*-like CNR in this species, the public assemblies were not sufficient to assess the genomic structure.

Regarding the intragenomic heterogeneity of the rDNA, the public genomes contained too many ambiguous sites to allow a discussion of the heterogeneity. Hence, we skipped the analysis of the intragenomic heterogeneity in the *non-Rhizophagus* group.

### Distribution of intragenomic polymorphisms is highly similar between the two *Rhizophagus* species

We then focused on the conservation of the intragenomic polymorphic site among the AM species. A previous study of *R. irregularis* reported that the intragenomic polymorphic site is not distributed uniformly on the rDNAs and argued that the variation-enriched site may be upon the diversifying selection [5]. However, a report of a single species is insufficient to exclude the possibility of the accumulation of random mutations at that site. Hence, we compared the intragenomic polymorphic site between the two *Rhizophagus* species, and found a similarity of the polymorphic site distribution. The variation-enriched site corresponded with the yeast ES27L, which is one of the eukaryote-specific (ES) expansion segments related to translation fidelity and antibiotic sensitivity [26–28].

To quantify the intragenomic polymorphism, we calculated the intragenomic polymorphic sites for each 50-base window on the aligned rDNA copies. As references, we selected *C. merolae* and a malaria parasite species, *Plasmodium falciparum,* due to the similarity of their rDNA structure with that of AM fungi [29, 30]. The overview of the polymorphism distribution was highly similar between *R. irregularis* and *R. clarus*, although the remaining two species indicated several different patterns with them (**Fig. 3**). Many of the variations were located on the ITS regions in all samples. The AM species and *P. falciparum* had an additional peak on the forward region of 28S. The two AM fungi shared another peak in the middle of the 28S rDNA. We then identified the corresponding region with the six domains on the *S. cerevisiae* 28S rDNA [31] via alignment, and revealed that the variation-enriched region in the AM fungi corresponded with domains I, III, IV, and VI. Intriguingly, the ES27L site of domain IV exhibited an especially high variation in both species (**Fig. 3**).

**Fig. 3.**
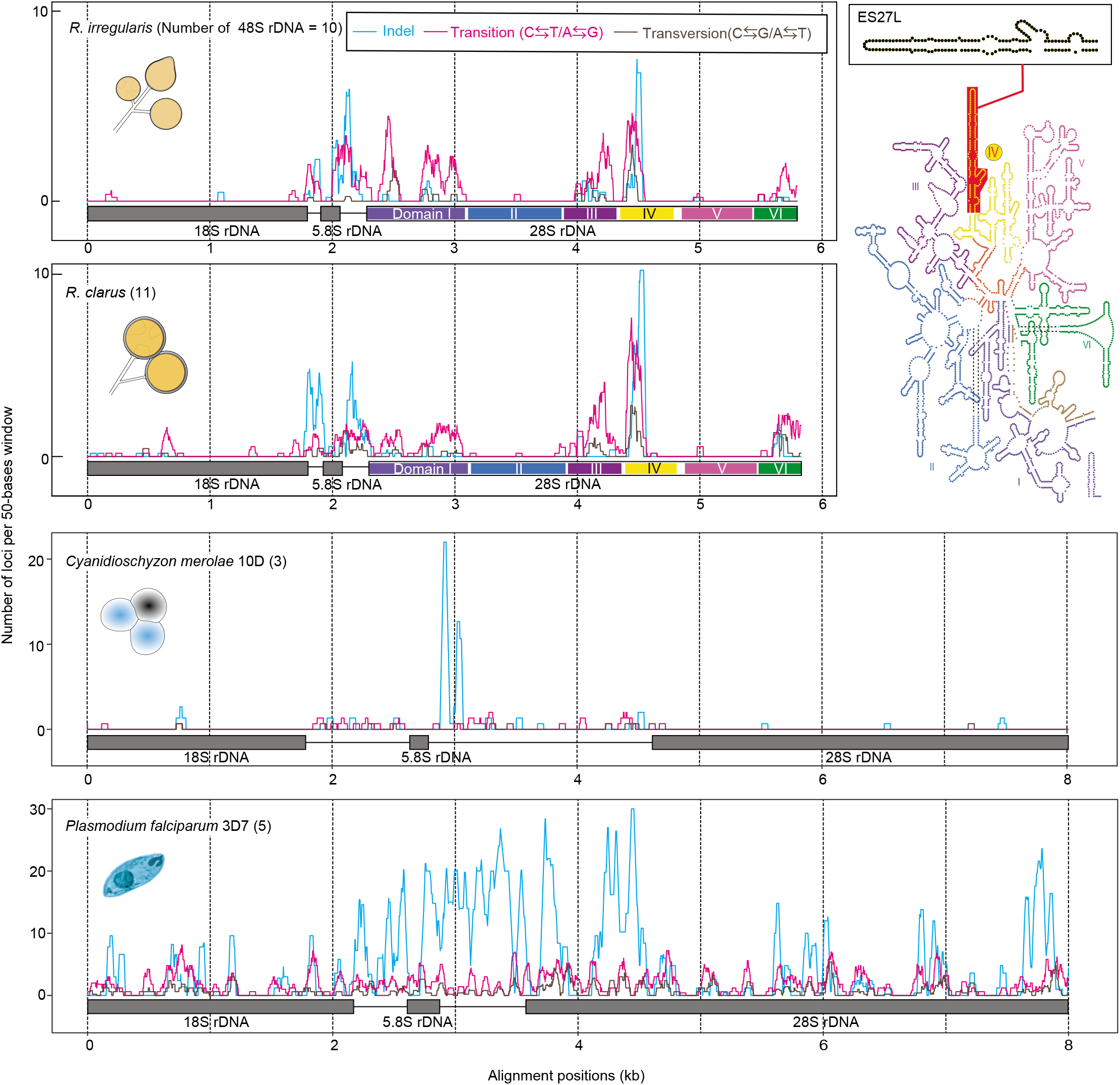
Distribution of rDNA sequence variants within the 48S rDNA. Numbers of 48S rDNA in each species were described next to the right of the species name. The boxes above the x-axis indicate the 18S-5.8S-28S ribosomal RNA-encoding region. The secondary structure of the *S. cerevisiae* 28S rRNA was placed on the top right. The structure corresponding to ES27L was highlighted in red and magnified in the sub-box.

To assess the effect of the polymorphism on the rRNA secondary structure, we performed an *in silico* structure prediction. Before the prediction, we clustered the rDNA copies that had identical “domain IV” sequences and obtained five *R. irregularis* clusters and four *R. clarus* clusters (**Fig. 4a**). Our *in silico* analysis predicted that the AM fungi have three types of the “ES27L” structure; i.e., a *S. cerevisiae*-like structure (named yeast type), a “c” arm-lacking structure (straight type), and a structure with an additional branch on the “b” arm (beak type) (Fig. 4B). Although the beak type was only found in *R. clarus* (c3086-1), the yeast type and straight type were detected in the rDNA of both species. The yeast type was abundant in *R. irregularis* (eight in *R. irregularis* and two in *R. clarus*), and the straight type was more frequent in *R. clarus* (eight in *R. clarus* and two in *R. irregularis*). The phylogenetic analysis of the whole 28S rDNA sequences revealed that the copies that had the same structural type did not establish a monophyletic clade (**Fig. 4b**).

**Fig. 4.**
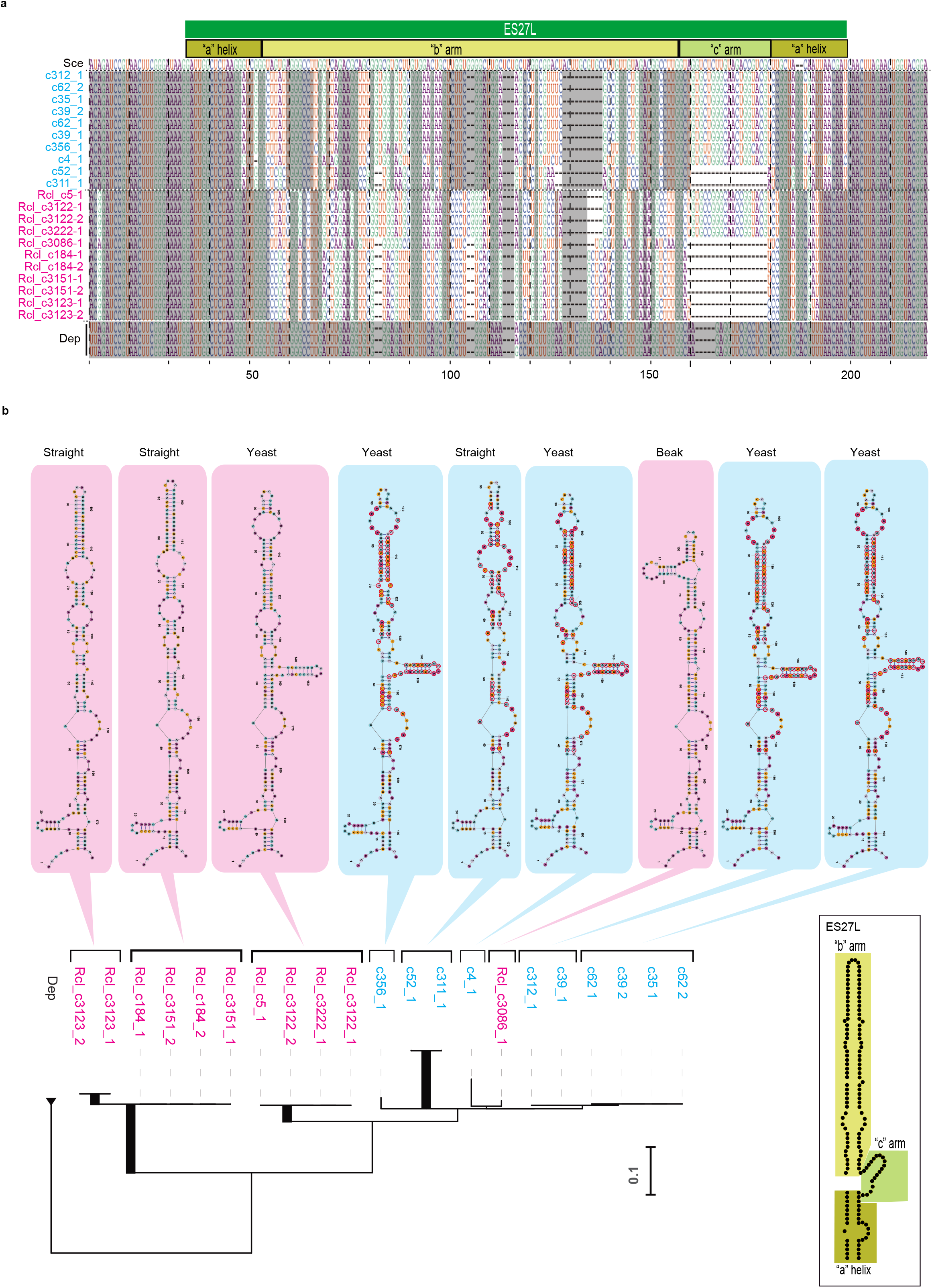
Polymorphisms on the ES27L region of the 28S rDNA/RNA. **a** Alignment of ES27L regions. The intragenomic conserved positions within each species were masked with gray color. **b** Secondary structures obtained from the *in silico* prediction of domain IV. Magenta, *R. clarus*; cyan, *R. irregularis*. The bottom ML tree was prepared from the whole 28S rDNA sequence. The abbreviations are defined in Supplementary Data 2.

### Host-specific rDNA expression in *R. irregularis*

To examine whether the rDNA expression profiles were affected by host conditions or the host species, we performed a PacBio CCS, which generates accurate long-read sequences through multiple repetitions of the sequencing of the same DNA molecules [19]. This method has been adopted in several ecological studies targeting rDNA in AM species [32, 33]. Our rRNA-targeting CCS showed that the rDNA expression profiles largely depended on the host plant species.

We constructed 23 CCS libraries from *R. irregularis*-colonizing plant organs (**Supplementary Data 4) (INSDC; PRJDB9672**). As a host plant, we used the model legume *Lotus japonicus* MG20 [34, 35] (17 libraries) and the basal land plant *Marchantia paleacea* [36] (6 libraries). We cultivated the legume-infecting samples under five conditions (**Supplementary Data 4**) to analyze the expression profiles after cuttings off the host shoots. The shoots cutting promotes the sporulation of AM fungi [37]. We obtained 61,290 AM fugal reads from the libraries and clustered the CCS reads to calculated the relative amount of each rRNA type. After several rounds of optimization of the CD-Hit-est-2D [38] parameters, we obtained the best clustering of the sequences with -c 99.

The identified profiles showed a conserved rRNA expression ratio in *L. japonicus* mycorrhizal roots with or without the shoot-cutting step in the host plants. Conversely, we detected significant differences in their expression profiles between *L. japonicus* roots and *M. paleacea* thalli (**Fig. 5**). The trend of the profiles was similar in types [c62_1, c62_2], [c312_1], [c356_1], and [c4_1]. However, it was different in the remaining three types ([c35_1], [c39_1], and [c52_1, c311_1]), which exhibited different rankings in the degree of expression according to the cultivation condition (**Fig. 5A**). The PCA analysis of the rRNA profiles revealed that samples of colonized *L. japonicus* roots were indistinguishable under the various conditions and that the clear distinction of profiles depended on the host species (**Fig. 5b**). The multivariate analysis of variance (MANOVA) identified statistically significant differences in the profiles *(P* < 0.001) (**Fig. 5b**).

**Fig. 5.**
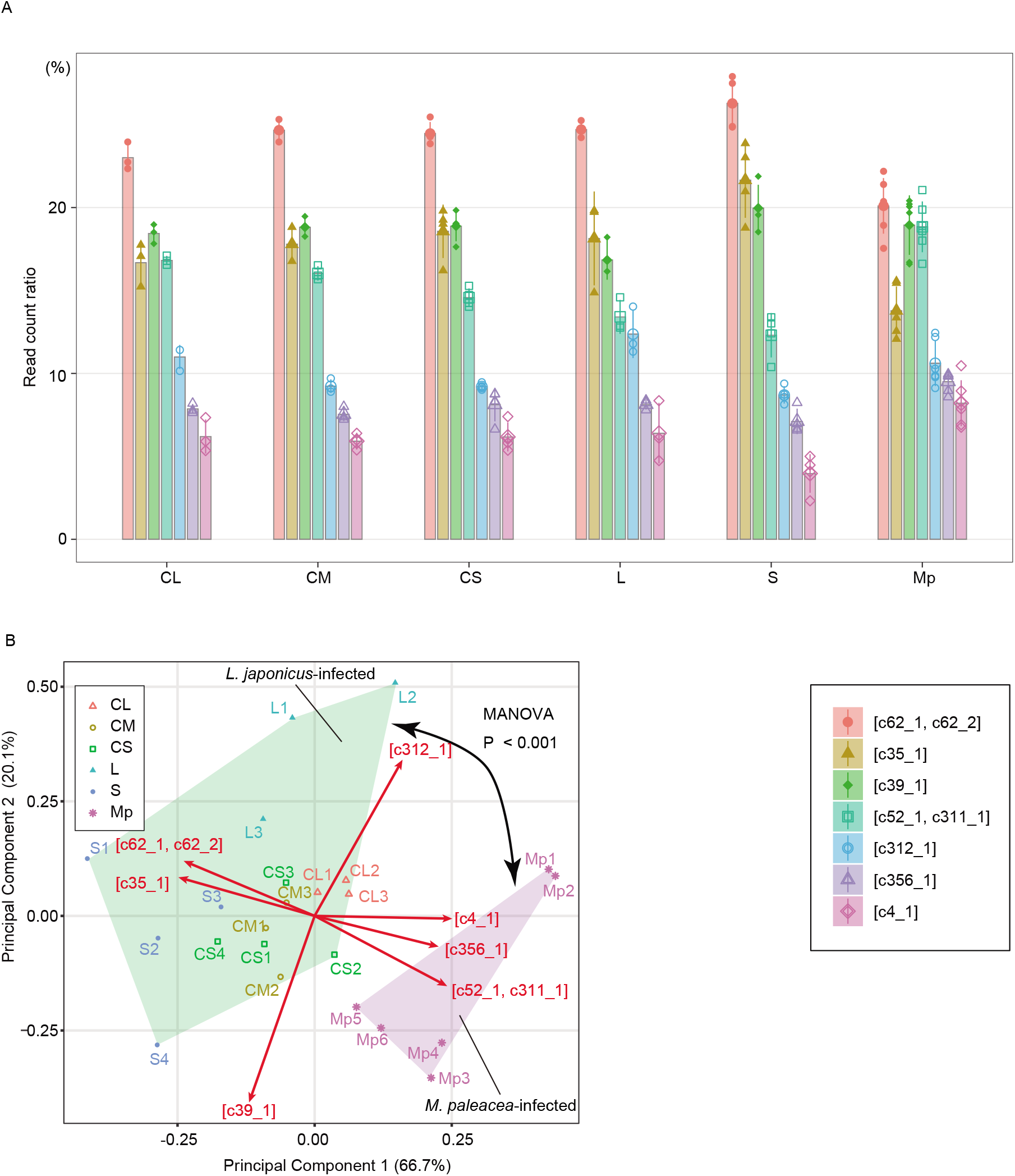
Expression profiles of the rDNA in *R. irregularis* colonizing different host species and at different growth conditions. **a** Read count ratio in each AM fungal CCS library. Each point indicates the value from a sample (error bar = standard deviation). The bar represents the average of the values. The colors and point shapes indicate the type of rDNA. Legends are provided in the right-bottom box. The x-axis represents the condition of the colonized root samples of *L. japonicus* MG20 (CL-S). CL, shoot cutting at 22 days postinoculation (dpi) and sampling at 36 dpi; CM, shoot cutting at 22 dpi and sampling at 28 dpi; CS, shoot cutting at 22 dpi and sampling at 24 dpi; L, cultivated for 36 dpi; S, cultivated for 24 dpi; MP, inoculated in *M. paleacea* and cultivated for three months. **b** Biplot of the principal component analysis (PCA) and their principal component score (red arrow). Each point represent a sample. The shape and color of the points indicate the difference of host species and growth conditions (see the top left box). Sample names were described to near the points.

## Discussion

Here, we revealed that the three rDNA features of *R. irregularis* were conserved in *R. clarus* [5, 14], i.e., (1) decreased copy count of the 48S rDNA, (2) absence of the TRS, and (3) intragenomic heterogeneity. Moreover, our analysis of the public genome data supported the wide distribution of the disarranged rDNA (a handful of TRS-lacking rDNAs) in non-*Rhizophagus* AM fungi. The cross-species comparison found conservation of the variation-enriched sites, ES27L, between *R. irregularis* and *R. clarus*. The rRNA-targeting clustering showed that the rRNA expression profiles were affected by the host plant species.

Based on the improved/reanalyzed genome data, we found that three AM species commonly had an exceptionally low 48S rDNA CN *(R. clarus,* 11 copies; *G. cerebriforme,* 2 copies; and *D. epigaea,* 7–10 copies). The previously reported CN in *R. irregularis* (ten copies) was the lowest among eukaryotes other than pneumonia-causing *Pneumocystis* (one copy) [39], *C. merolae* (three copies) [29], and malaria-causing *Plasmodium* (five to eight copies) [5, 30]. The CNV of the rDNA ranged from 14 to 1,442 in other fungi [23]. The rDNA CN is relevant for translation efficiency, because multiple rDNAs are required to synthesize a sufficient amount of rRNA [24, 40]. Although an rDNA CN <20 is lethal in *S. cerevisiae* [24], the successive cultivation of AM fungi with their reduced rDNA counts suggested that the handful of rDNA copies is sufficient to support the growth of AM fungi. Our team previously hypothesized that the AM fungus has a unique ribosome synthesis step to recover from the low rDNA CN, based on the results obtained for a single AM species, *R. irregularis* [5]. The conservation of the rDNA CNR identified here supports the generalization of our previous hypothesis to the genus *Rhizophagus* and the AM fungus.

Our depth-based CNV analysis of non-AM fungi suggested that a sister group of the AM fungus, *Mucorales,* retained the ordinary number of rDNA copies (71–151) (**Fig. 2, Supplementary Data 2**). The simple mapping to the phylogenetic tree indicated that the CNR occurred on the common ancestor of the AM fungus, although rDNA data for *Endogonales* (a plant-symbiotic group closely related to *Mucorales)* [25] have not been collected (**Supplementary Figure 2**). It is interesting that a previous study identified CNR on distantly related plant-symbiotic fungi (e.g., *Oidiodendron maius,* 11 copies; *Phialocephala scopiformis*, 15 copies; *Cenococcum geophilum*, 15 copies; and *Meliniomyces variabilis*, 18 copies) [23]. Although the genomic structure and intragenomic heterogeneity of these species remain unknown, the rDNA CNR may be a universal trend related to plant root symbiosis.

We also found evidence of the conservation of the absence of the TRS in AM fungi. The TRS-lacking feature has been observed only in the *R. irregularis*, malaria parasites, and *C. merolae* genomes, which exhibit 48S rDNA CNR (malaria parasites, five to eight copies; *C. merolae*, three copies). This correspondence between the TRS-lacking and CNR features is reasonable because the TRS is part of the recovery system of the rDNA CN, i.e., unequal sister chromatid recombination (USCR) [41]. The TRS increases the rDNA CN via recombination with a neighbor repeat during the DNA duplication stages [42]. An unknown highly efficient rRNA synthesis activity in the AM fungus might reduce the selective pressure against the retention of the rDNA CN and might disrupt the TRS in AM fungi. We found both TRS-lacking and TRS-keeping 48S rDNAs in the two *Mucorales* species (**Fig. 2**). Although the genomic structure of the remaining rDNA copies in these species is unclear, the TRS-lacking rDNA might have arisen from a normal TRS-containing rDNA in the common ancestor of the AM fungus and *Mucorales*; subsequently, the ancestral AM fungus might have lost the TRA-containing rDNAs.

The heterogeneity of the intragenomic sequence in rDNA is attractive in terms of ribosome heterogeneity. The evolutional model of the rDNA has assumed that the TRS-dependent CN recovery causes the homogeneity of the rDNA through the bottleneck effect (concerted evolution) [24]. Several organisms, including the TRS-lacking malaria parasites, exhibit rDNA/rRNA heterogeneity with tissue- and cell stage-specific expression patterns [43–45]. However, the functional consequences of this rRNA variation have not been established [2], and It was also unclear whether the rDNA heterogeneity is related with an adaptation or an evolutionarily neutral diversification. We revealed the conservation of polymorphism-enriched region of ITS, domains I, III, IV, and VI, between the two *Rhizophagus* (**Fig. 3**). For the ITS region, a similar variation accumulation was determined from other TRS-lacking species (*C. merolae* and *P. falciparum*). This accumulation may be attributed to their weak evolutional constraints compared with rRNA-encoding regions. Conversely, the variation peak on the ES27L site of domain IV was only found on the two AM fungi; the polymorphism of *C. merolae* was very weak on the rRNA-encoded region, and *P. falciparum* exhibited a scattered distribution of the variation site for all RNA-encoded regions (**Fig. 3**). The data concerning *C. merolae* indicate that the polymorphism on ES27L is not inevitable in TRS-lacking eukaryotes. These results suggest that the polymorphisms did not occur randomly on the AM rDNAs, but they reflected some functional redundancy or disruptive selection on the region. Specific alleles on this site may be maintained for unknown evolutional reasons.

Our *in silico* secondary structure analysis indicated that intragenomic variations were scattered over the “b” stem loops of *Rhizophagus* ES27L (**Fig. 4**). Moreover, the two AM fungi commonly had an rDNA genotype lacking the “c” arm (straight type). In multiple species, the ES27L has been modeled in cryo-EM, to point toward the peptide exit tunnel of the ribosome [46, 47]. A previous experimental deletion indicated that the “b” arm has a function in translation fidelity via binding to methionine aminopeptidase (MAP1) in *S. cerevisiae*; the deletions in the “b” arm induce amino acid misincorporation and stop codon read through upon treatment with translational-error-inducing antibiotics (paromomycin) [27], and the whole deletion of “b” changes the proteomic profiles [28]. The polymorphism observed on the AM fungal “b” arm may contribute to translation fidelity control and modulates the sensitivity against antibiotics. The heterogeneity observed in ES27L was not accompanied by the diversification of the binding protein on the platform; two *Rhizophagus* genomes contained the single ortholog of yeast Map1, respectively (Supplementary Data 6). It should be noted that the deletions in the “b” arm increased the resistance against anisomycin and cycloheximide in yeast [27], indicating that the complete “b” arm reduces the resistance against these antibiotics. This trade-off might result in the selection of the diversification of the AM fungal rDNA. It should be noted that plants have an antibiotic-like substrate that is used for self-defense [48]. For example, ricin from *Ricinus communis* cleaves the *N*-glycosidic bond in 28S rRNA [49]. The intragenomic diversity of rDNA may enable symbionts to pass through the species-specific diversified defense mechanisms of plants and contribute to the establishment of symbiosis with a broad range of plant species.

Regarding the “c” arm, the deleted yeast did not display any changes in the sensitivity or tolerance to any of the inhibitors tested (cycloheximide, anisomycin, and hygromycin B) [27]. However, the conservation of the “c” arm among eukaryotes suggests their importance for life. The deletion of the whole ES27L is lethal in yeast and *Tetrahymena* [26, 50]. Meanwhile, in *Drosophila* ribosomes, the interaction between the “c arm” and the S8e ribosomal protein forms an intersubunit bridge and was considered to contribute to the conformation dynamics of the binding with the elongation factor 2 (eEF2) [51]. Nonetheless, future studies of the effect against various antibiotics are needed to reveal the function of ES27L in AM species.

The heterogeneity of the rDNA is also an important subject in terms of biodiversity. Our *R. clarus* genome assembly indicated that intragenomic variation covered most of the polymorphisms previously reported in this species (**Fig. 1d**). This result provides an incentive to review the rDNA diversity of other AM fungi. The ITS and a part of the 28S rDNA have been widely used as a marker gene for the ecological and taxonomic studies of AM fungi [52]. The intraindividual rDNA diversity of AM fungi has been indicated because multiple rDNA genotypes were reported from the single fungal body in many AM species [53–57]. Singlenucleus sequencing indicated that this multinucleate fungus has genetically different nuclei in its body (heterokaryosis) [14, 15, 58]. Our previous finding of intragenomic heterogeneity in *R. irregularis* led to a discussion of the contribution of the combination of heterokaryosis and intragenomic heterogeneity to intraindividual heterogeneity within a fungal body [5]. The present results from *R. clarus* reinforce the hypothesis that the multilayered diversification mechanism causes intraindividual rDNA heterogeneity in *Rhizophagus*. The intragenomic variation in *R. irregularis* was not sufficient to disrupt species-level identification [5]. However, our result that an *R. clarus* rDNA copy established a clade with the sequence of *R. cactus* (**Fig. 1d**) indicated that imbalanced amplification and sequencing among the paralogs have the potential to cause erroneous identification of species.

The rRNA profile of *R. irregularis* identified here was similar under the same incubation condition but was modified by drastic changes in the condition. A previous RNA-seq study indicated that all of the rDNA genotypes retained the translation activity in a pre-infected stage of *R. irregularis* [5]. The rRNA profiles were significantly affected by the variation of the incubation period (22–36 dpi and 3 months) and the type of host plant (legumes to liverworts). Moreover, the ratio of each rRNA type was not correlated with the CN of each rDNA gene (**Fig. 5**). Due to the slow growth rate of liverwort-infected AM fungi, we cannot align the incubation period with that of the legume-infected sample. However, the observed dynamics suggest that the AM fungus changed the 28S rDNA expression of each copy via a yet unknown environment-specific expression control system. Other species containing heterogeneous rDNA/rRNA (malaria parasites, zebrafish, mice, and humans) [43, 44, 51, 59] also exhibit similar condition-dependent expression changes. Although these organisms have complete replacements of the rRNA type at some developmental stages [59], we found no replacement of the rRNA types under the adopted conditions. We used the whole body of the incubated AM fungi together with host plant roots. Additional structure-specific rRNA sampling would provide insights into the dynamics and their adaptive contribution to AM fungi. The principal component score and the actual change in the reading count ratio indicated that the change of [c52_1, c311_1]-type expression ratio relates to the host-dependent modification. Interestingly, the [c52_1, c311_1] type lacks the “c” arm on the ES27L, and some SNP on “b” arms were determined based on a comparison with the remaining types (**Fig. 4**). This result tempted us to speculate that this [c52_1, c311_1]-derived rRNA affords a suitable ribosome for the symbiosis with liverworts.

Here, we indicated the possibility of relationships between plant symbiosis and disarranged heterogeneous rDNA. Although multiple genomes of mutualistic eukaryote have been identified [60]. Extended genomes: symbiosis and evolution), the previous studies ignored the analysis of rDNA due to the difficulty in assembling their genome. The assumption that all the eukaryotes have homogeneous TRS-making rDNAs may have drawn attention away from rDNA diversity. The recent “renaissance” of ribosome heterogeneity [2] may renew the study not only of embryology/physiology but also of symbiotic biology.

## Material and Methods

### PacBio-based genome assembling

The genome sequencing and gene annotation of *R. clarus* HR1 (MAFF:520076) was performed according to a previous study [5], with some modification (Supplementary Data 9). We isolated a complete mitochondrial sequence from the contigs and then submitted the assemblies to the DDBJ (nuclear DNA = BLAL01, mitochondrial DNA = LC506577). For the prompt submission of the genomic gene model data to DDBJ, we used GFF2MSS ver. 3.0.2 script (https://github.com/maedat/GFF2MSS).

### Detection of ribosomal DNA and intragenomic polymorphisms

Ribosomal DNA regions were detected by RNAmmer ver. 1.284 [61]. In *R. clarus,* the RNAmmer-based regions were refined manually based on the MAFFT v7.429-based [62] alignment to the 48S rRNA of *S. cerevisiae* S288C. The genomic positions of rDNAs were visualized using our script, GeneHere version 0.1.1 (https://github.com/maedat/GeneHere). The species analyzed and genomic data are summarized in **Supplementary Data 2**. We adopted the rDNA searches against 26 species and obtained multiple 18S and 28S sequences from 12 species. We excluded the remaining samples containing only a part of the 48S rDNA sequence from the downstream analysis.

The number of rDNA paralogs was estimated based on the mean depth of coverage. From public short-read data, we found suitable data for nine species (**Supplementary Data 2**). The obtained read data were mapped to the references (whole-genomic data, extracted 18S, or 28S rDNA sequences) using bowtie2 version 2.3.5.1. The coverage depth of the references was calculated using bedtools version v2.29.0 (bedtools genomecov command with -d option), and the statistics of each region were calculated and visualized using the R software ver. 3.6.1 with the ggplot2 ver. 3.2.1 package.

The difference among the rDNA paralogs was calculated using our script, AliVa (https://github.com/maedat/AliVa), and the sequences were aligned by MAFFT ver. 7.427 (options: --auto). The neighbor-net tree presented in **Fig. 1b** was generated by SplitsTree4 ver. 4.14.8 (raw sequence data: 10.6084/m9.figshare.11880780) [63]. The phylogenetic trees depicted in Fig. 1c were constructed based on the MAFFT alignment (10.6084/m9.figshare.11880834) using the ML method with IQ-TREE ver.1.6.11 (options: -nt AUTO) [64]; they were also tested for robustness by bootstrapping (1000 pseudoreplicates).

### Analysis of the secondary structure of rDNA

The secondary structure of the *Rhizophagus* 28S rDNAs was predicted in silico. Domains I-VI were determined through the manual alignment with the *S. cerevisiae* 28S rDNA. We then cut out each domain from the *Rhizophagus* rDNAs and predicted the secondary structure of each domain. We used RNAstructure ver. 6.1 for structure prediction and StructureEditor ver 6.1 for visualization [65]. After the prediction of the top five minimum free energy structures, we chose the structure that exhibited the greatest similarity to that of *S. cerevisiae*.

### Expression dynamics of rRNA

The expression levels of the rDNA paralogs were examined with PacBio sequel in total RNA extracted from plant-infected *R. irregularis* DAOM-181602. Thalli of *Marchantia paleacea* subsp. *diptera* were grown with *R. irregularis* DAOM-181602 (Premier Tech) in growth conditions, as described previously [36].

Total RNAs were isolated using the modified CTAB method, as described previously by Nakagawa et al. (2011) [66], with some modifications (Supplementary Data 9). *Lotus japonicus* MG-20 seeds were surface sterilized and germinated on an agar plate containing no nutrients. Plants were grown in an artificially lit growth cabinet at 24°C for 16 h (light) and 22°C for 8 h (dark). After 6 days, the seedlings were transferred to soil, as described previously by Miyata et al. (2014), with or without spores of *R. irregularis.* After 22 days, some of these plants were cut, their aerial parts were removed, and plants were harvested at 2, 6, or 14 days after shoot removal.

The library used for PacBio CCS was prepared according to the protocol “Full-Length 16S Amplification, SMRTbell Library Preparation and Sequencing” (Pacific Biosciences, Part Number 101-599-700 version 01), with some modifications (Supplementary Data 9). The CCS sequences were generated on a PacBio Sequel sequencer using a Sequel Binding and Internal Control Kit 3.0 Mag Bead Binding Buffer Kit v2, and Sequel Sequencing Kit 3.0 (Pacific Biosciences). The raw reads obtained were assembled using SMRTLINK7 (Pacific Biosciences).

## Author contributions

TM and MK conceived of and designed the experiments; TM, YK, TE, KY, TB, YN, SS, and MK performed the *R. clarus* genomic experiments and analyses; TM, TN, KY, SS, and MK performed the *R. irregularis* transcriptomic experiments and analyses. TM, TN, TE, SS, MK wrote the paper.

## Acknowledgment

This work was supported by JST ACCEL Grant Number JPMJAC1403 and MEXT/JSPS KAKENHI Grant Number 19K22269, Japan. We thank Miwako Matsumoto, the Functional Genomics Facility and the Data Integration and Analysis Facility at the National Institute for Basic Biology for technical support; Katsuharu Saito, Kohki Akiyama. Computations were partially performed on the NIG supercomputer at ROIS National Institute of Genetics.

## References

1. Crick FH. On protein synthesis. Symp Soc Exp Biol 1958; 12: 138–163.

2. Genuth NR, Barna M. The discovery of ribosome heterogeneity and its implications for gene regulation and organismal life. Mol Cell 2018; 71: 364–374.

3. Dinman JD. Pathways to specialized ribosomes: the brussels lecture. J Mol Biol 2016; 428: 2186–2194.

4. Xue SF, Barna M. Specialized ribosomes: a new frontier in gene regulation and organismal biology. Nat Rev Mol Cell Biol 2012; 13: 355–369.

5. Maeda T, Kobayashi Y, Kameoka H, Okuma N, Takeda N, Yamaguchi K, et al. Evidence of non-tandemly repeated rDNAs and their intragenomic heterogeneity in *Rhizophagus irregularis*. Commun Biol 2018; 1: 87.

6. Spatafora JW, Chang Y, Benny GL, Lazarus K, Smith ME, Berbee ML, et al. A phylum-level phylogenetic classification of zygomycete fungi based on genome-scale data. Mycologia 2016; 108:1028–1046.

7. Bonfante P, Genre A. Mechanisms underlying beneficial plant–fungus interactions in mycorrhizal symbiosis. Nat Commun 2010; 1: 48.

8. Corradi N, Bonfante P. The arbuscular mycorrhizal symbiosis: origin and evolution of a beneficial plant infection. PLoS Pathog 2012; 8.

9. Öpik M, Davison J. Uniting species-and community-oriented approaches to understand arbuscular mycorrhizal fungal diversity. Fungal Ecology. 2016., 24: 106–113

10. van der Heijden MGA, Klironomos JN, Ursic M, Moutoglis P, Streitwolf-Engel R, Boller T, et al. Mycorrhizal fungal diversity determines plant biodiversity, ecosystem variability and productivity. Nature 1998; 396: 69–72.

11. Van Der Heijden MGA, Martin FM, Selosse M-A, Sanders IR. Mycorrhizal ecology and evolution: the past, the present, and the future. New Phytol 2015; 205: 1406–1423.

12. Davison J, Moora M, Ouml;pik M, Adholeya A, Ainsaar L, Ba A, et al. Global assessment of arbuscular mycorrhizal fungus diversity reveals very low endemism. Science 2015; 349: 970–973.

13. Gianinazzi-Pearson V. Plant cell responses to arbuscular mycorrhizal fungi: getting to the roots of the symbiosis. Plant Cell 1996; 8: 1871–1883.

14. Lin K, Limpens E, Zhang Z, Ivanov S, Saunders DGO, Mu D, et al. Single nucleus genome sequencing reveals high similarity among nuclei of an endomycorrhizal fungus. PLoS Genet 2014; 10: e1004078.

15. Montoliu-Nerin M, Sánchez-García M, Bergin C, Grabherr M, Ellis B, Kutschera VE, et al. Building *de novo* reference genome assemblies of complex eukaryotic microorganisms from single nuclei. Sci Rep 2020; 10: 1303.

16. Morin E, Miyauchi S, San Clemente H, Chen ECH, Pelin A, de la Providencia I, et al. Comparative genomics of *Rhizophagus irregularis, R. cerebriforme, R. diaphanus* and *Gigaspora rosea* highlights specific genetic features in Glomeromycotina. New Phytol 2019; 222:1584–1598.

17. Venice F, Ghignone S, Salvioli di Fossalunga A, Amselem J, Novero M, Xianan X, et al. At the nexus of three kingdoms: the genome of the mycorrhizal fungus *Gigaspora margarita* provides insights into plant, endobacterial and fungal interactions. Environ Microbiol 2020; 22: 122–141.

18. Kobayashi Y, Maeda T, Yamaguchi K, Kameoka H, Tanaka S, Ezawa T, et al. The genome of *Rhizophagus clarus* HR1 reveals a common genetic basis for auxotrophy among arbuscular mycorrhizal fungi. BMC Genomics 2018; 19: 465.

19. Wenger AM, Peluso P, Rowell WJ, Chang P-C, Hall RJ, Concepcion GT, et al. Accurate circular consensus long-read sequencing improves variant detection and assembly of a human genome. Nat Biotechnol 2019; 37: 1155–1162.

20. Cohn M, Liti G, Barton DBH. Telomeres in fungi. Comparative Genomics. 2005. Topics in current genetics, pp 101–130.

21. Kamel L, Keller-Pearson M, Roux C, Ané J-M. Biology and evolution of arbuscular mycorrhizal symbiosis in the light of genomics. New Phytol 2017; 213: 531–536.

22. Walker BJ, Abeel T, Shea T, Priest M, Abouelliel A, Sakthikumar S, et al. Pilon: an integrated tool for comprehensive microbial variant detection and genome assembly improvement. PLoS One 2014; 9:e112963.

23. Lofgren LA, Uehling JK, Branco S, Bruns TD, Martin F, Kennedy PG. Genome based estimates of fungal rDNA copy number variation across phylogenetic scales and ecological lifestyles. Mol Ecol 2019; 28: 721–730.

24. Ganley ARD, Kobayashi T. Highly efficient concerted evolution in the ribosomal DNA repeats: Total rDNA repeat variation revealed by whole-genome shotgun sequence data. Genome Res 2007; 17: 184–191.

25. Chang Y, Desirò A, Na H, Sandor L, Lipzen A, Clum A, et al. Phylogenomics of Endogonaceae and evolution of mycorrhizas within Mucoromycota. New Phytol 2019; 222: 511–525.

26. Ramesh M, Woolford JL Jr. Eukaryote-specific rRNA expansion segments function in ribosome biogenesis. RNA 2016; 22: 1153–1162.

27. Fujii K, Susanto TT, Saurabh S, Barna M. Decoding the function of expansion segments in ribosomes. Mol Cell 2018; 72: 1013–1020.e6.

28. Shankar V, Rauscher R, Reuther J, Gharib WH, Koch M, Polacek N. rRNA expansion segment 27Lb modulates the factor recruitment capacity of the yeast ribosome and shapes the proteome. Nucleic Acids Res 2020.

29. Matsuzaki M, Misumi O, Shin-I T, Maruyama S, Takahara M, Miyagishima S-Y, et al. Genome sequence of the ultrasmall unicellular red alga *Cyanidioschyzon merolae* 10D. Nature 2004; 428: 653–657.

30. Gardner MJ, Hall N, Fung E, White O, Berriman M, Hyman RW, et al. Genome sequence of the human malaria parasite Plasmodium falciparum. Nature 2002; 419: 498–511.

31. Petrov AS, Bernier CR, Gulen B, Waterbury CC, Hershkovits E, Hsiao C, et al. Secondary structures of rRNAs from all three domains of life. PLoS One 2014; 9: e88222.

32. Schlaeppi K, Bender SF, Mascher F, Russo G, Patrignani A, Camenzind T, et al. High-resolution community profiling of arbuscular mycorrhizal fungi. New Phytol 2016; 212: 780–791.

33. Tedersoo L, Tooming-Klunderud A, Anslan S. PacBio metabarcoding of fungi and other eukaryotes: errors, biases and perspectives. New Phytol 2018; 217: 1370–1385.

34. Kawaguchi M. *Lotus japonicus* ‘miyakojima’ mg-20: an early-flowering accession suitable for indoor handling. J Plant Res 2000; 113: 507–509.

35. Kato T, Kaneko T, Sato S, Nakamura Y, Tabata S. Complete structure of the chloroplast genome of a legume, *Lotus japonicus*. DNA Res 2000; 7: 323–330.

36. Kobae Y, Ohtomo R, Morimoto S, Sato D, Nakagawa T, Oka N, et al. Isolation of native arbuscular mycorrhizal fungi within young thalli of the liverwort *Marchantia paleacea*. Plants 2019; 8.

37. Redhead JF. Endotrophic mycorrhizas in Nigeria: some aspects of the ecology of the endotrophic mycorrhizal association of *Khaya grandifoliola* C. DC. Endomycorrhizas; Proceedings of a Symposium 1975.

38. Fu L, Niu B, Zhu Z, Wu S, Li W. CD-HIT: accelerated for clustering the next-generation sequencing data. Bioinformatics 2012; 28: 3150–3152.

39. Cushion MT, Keely SP. Assembly and annotation of *Pneumocystis jirovecii* from the human lung microbiome. MBio 2013; 4.

40. Milo R, Phillips R, Goodsell DS, Lane N, Nelson P, Hoffmann PM, et al. Cell biology by the numbers. Taylor & Francis Inc. https://www.bookdepository.com/Cell-Biology-by-Numbers-Ron-Milo/9780815345374. Accessed 5 Mar 2018.

41. Ganley ARD, Ide S, Saka K, Kobayashi T. The effect of replication initiation on gene amplification in the rDNA and its relationship to aging. Mol Cell 2009; 35: 683–693.

42. Kobayashi T. Ribosomal RNA gene repeats, their stability and cellular senescence. Proceedings of the Japan Academy Series B-Physical and Biological Sciences 2014; 90: 119–129.

43. Gunderson JH, Sogin ML, Wollett G, Hollingdale M, de la Cruz VF, Waters AP, et al. Structurally distinct, stage-specific ribosomes occur in *Plasmodium*. Science 1987; 238: 933–937.

44. Li J, Wirtz RA, McConkey GA, Sattabongkot J, McCutchan TF. Transition of *Plasmodium vivax* ribosome types corresponds to sporozoite differentiation in the mosquito. Mol Biochem Parasitol 1994;65:283–289.

45. Waters AP. The ribosomal RNA genes of *Plasmodium*. Adv Parasitol 1994; 34: 33–79.

46. Armache J-P, Jarasch A, Anger AM, Villa E, Becker T, Bhushan S, et al. Cryo-EM structure and rRNA model of a translating eukaryotic 80S ribosome at 5.5-A resolution. Proc Natl Acad Sci U SA 2010;107: 19748–19753.

47. Greber BJ, Boehringer D, Montellese C, Ban N. Cryo-EM structures of Arx1 and maturation factors Rei1 and Jjj1 bound to the 60S ribosomal subunit. Nat Struct Mol Biol 2012; 19: 1228–1233.

48. Savoia D. Plant-derived antimicrobial compounds: alternatives to antibiotics. Future Microbiol 2012; 7: 979–990.

49. Endo Y, Tsurugi K. The RNA N-glycosidase activity of ricin A-chain. The characteristics of the enzymatic activity of ricin A-chain with ribosomes and with rRNA. J Biol Chem 1988; 263: 8735–8739.

50. Sweeney R, Chen L, Yao MC. An rRNA variable region has an evolutionary conserved essential role despite sequence divergence. Mol Cell Biol 1994; 14: 4203–4215.

51. Parks MM, Kurylo CM, Dass RA, Bojmar L, Lyden D, Vincent CT, et al. Variant ribosomal RNA alleles are conserved and exhibit tissue-specific expression. Sci Adv 2018; 4: eaao0665.

52. Krishnamoorthy R, Premalatha N, Karthik M, Anandham R, Senthilkumar M, Gopal NO, et al. Molecular markers for the identification and diversity analysis of arbuscular mycorrhizal fungi (AMF). Fungal Biology. 2017., 177–199

53. Sanders IR, Alt M, Groppe K, Boller T, Wiemken A. Identification of ribosomal DNA polymorphisms among and within spores of the *Glomales*: application to studies on the genetic diversity of arbuscular mycorrhizal fungal communities. New Phytologist. 1995., 130: 419–427

54. LloydMacgilp SA, Chambers SM, Dodd JC, Fitter AH, Walker C, Young JPW. Diversity of the ribosomal internal transcribed spacers within and among isolates of *Glomus mosseae* and related mycorrhizal fungi. New Phytol 1996; 133: 103–111.

55. Hosny M, Hijri M, Passerieux E, Dulieu H. rDNA units are highly polymorphic in *Scutellospora castanea* (Glomales, Zygomycetes). Gene 1999; 226: 61–71.

56. Hijri M, Sanders IR. The arbuscular mycorrhizal fungus *Glomus intraradices* is haploid and has a small genome size in the lower limit of eukaryotes. Fungal Genet Biol 2004; 41: 253–261.

57. Pawlowska TE, Taylor JW. Organization of genetic variation in individuals of arbuscular mycorrhizal fungi. Nature 2004; 427: 733–737.

58. Chen ECH, Morin E, Beaudet D, Noel J, Yildirir G, Ndikumana S, et al. High intraspecific genome diversity in the model arbuscular mycorrhizal symbiont *Rhizophagus irregularis*. New Phytol 2018.

59. Locati MD, Pagano JFB, Girard G, Ensink WA, van Olst M, van Leeuwen S, et al. Expression of distinct maternal and somatic 5.8S, 18S, and 28S rRNA types during zebrafish development. RNA 2017; 23: 1188–1199.

60. Hurst GDD. Extended genomes: symbiosis and evolution. Interface Focus 2017; 7: 20170001.

61. Lagesen K, Hallin PF, Rødland E, Stærfeldt HH, Rognes T, Ussery DW. RNammer: consistent annotation of rRNA genes in genomic sequences. Nucleic Acids Res 2007; 35: 3100–3108.

62. Katoh K, Standley DM. MAFFT multiple sequence alignment software version 7: improvements in performance and usability. Mol Biol Evol 2013; 30: 772–780.

63. Huson DH, Bryant D. Application of phylogenetic networks in evolutionary studies. Mol Biol Evol 2006; 23: 254–267.

64. Nguyen L-T, Schmidt HA, von Haeseler A, Minh BQ. IQ-TREE: a fast and effective stochastic algorithm for estimating maximum-likelihood phylogenies. Mol Biol Evol 2015; 32: 268–274.

65. Reuter JS, Mathews DH. RNAstructure: software for RNA secondary structure prediction and analysis. BMC Bioinformatics 2010; 11: 129.

66. Nakagawa T, Kaku H, Shimoda Y, Sugiyama A, Shimamura M, Takanashi K, et al. From defense to symbiosis: limited alterations in the kinase domain of LysM receptor-like kinases are crucial for evolution of legume-*Rhizobium* symbiosis. Plant J 2011; 65: 169–180.

