## Supplementary Data 9 for "Conservation and host-specific expression of non-tandemly repeated heterogenous ribosome RNA gene in arbuscular mycorrhizal fungi"

**The details of the protocols**

<Genome sequencing and gene annotation of *R. clarus* HR1>

DNA preparation: The DNA sample for the PacBio sequencing was extracted from a laboratory-incubated *R. clarus* HR1 (MAFF:520076), according to a previous study [[1]](https://paperpile.com/c/bDgfmN/xi8n).

PacBio sequencing: Long-read sequences were generated using a PacBio Sequel sequencer (Pacific Biosciences, Menlo Park, CA, USA) with a DNA/Polymerase Binding Kit P6v2 (Pacific Biosciences) and a DNA Sequencing Reagent Kit 4.0 v2 (Pacific Biosciences). The library was prepared according to the 14- or 18-kb template preparation of the BluePippin™ Size-Selection System (Sage Science, MA, USA). A total sequence of 19.7 Gb in 5,819,346 reads (134× coverage of the genome, assuming a genome size of 147 Mbp) was obtained from 21 SMRT cells (Supplementary Table 2). The N50 length of the raw reads was 13,107 bases.

PacBio-based genome assembly: The *R. clarus* genome was assembled using the CANU ver. 1.7.1 software [[2]](https://paperpile.com/c/bDgfmN/i9vf). We selected high-quality reads from the PacBio read (11 Gbp, 73× coverage of the genome) and then assembled the genome with the “genomeSize=150m” option. CANU was used to construct 469 contigs (147 Mbp). The obtained contigs were then polished with the PacBio read using ARROW ver. 2.2.2 [[3]](https://paperpile.com/c/bDgfmN/PnvB). For read mapping, we used bax2bam and pbalign (option: NPROC=48) and polished the aligned data with ARROW (option: NPROC=48).

Illumina-based error correction: After the polishing, erroneous regions were corrected with Illumina read (DRR121633 and DRR121632) using Pilon ver. 1.23 [[4]](https://paperpile.com/c/bDgfmN/6xFx). After the trimming of low-quality and adapter sequences using trim_galore ver. 0.5.0 (option: --trim1 --paired) (https://github.com/FelixKrueger/TrimGalore), Illumina reads were mapped with bowtie2 ver. 2.3.5.1 [[5]](https://paperpile.com/c/bDgfmN/LqPF). Pilon was then adopted with the default setting.

Gene prediction and annotation: The obtained assemblies were processed through a RNA-seq-based gene model construction pipeline, BRAKER-2.1.2 (Augustus version 3.3.2, GeneMark-ES Suite 4.32) [[6]](https://paperpile.com/c/bDgfmN/oAUD). As the referential data for gene model construction, we obtained standard RNA-seq data from INSDC (a total of 102 Gb, Supplementary Data 5). After filtering low-quality and adapter sequences with trim_galore ver. 0.5.0, RNA-seq data were then mapped to the genomic assemblies using hisat ver. 2.1.0 [[7]](https://paperpile.com/c/bDgfmN/P7TZ) with the default setting. As references, previously constructed gene models (GBB83119, GBC10880) from a draft *R. clarus* HR1 genome (PRJDB6444) were also aligned to the genome using exonerate ver. 2.2.0 [[8]](https://paperpile.com/c/bDgfmN/2OP8) (option --model est2genome --bestn 1 --percent 80). To train the probabilistic models of Augustus, we used WebAUGUSTUS [[9]](https://paperpile.com/c/bDgfmN/fhpw) with the referential data sets. After the soft masking of repetitive regions on the genome by redmask.py v0.0.2 (https://github.com/nextgenusfs/redmask), we adopted BRAKER with the referential data set (option --skipAllTraining --softmasking) and trained probabilistic models. As a result, we obtained 31,233 protein-encoding genes and 32,541 isoforms (Supplementary Data 6). The completeness of the constructed gene model was evaluated using BUSCO ver. 3.0.2 [[10]](https://paperpile.com/c/bDgfmN/1Kyc). The BUSCO analysis used the Fungi odb9 and Eukaryota odb9 as a benchmark gene set and employed the “-m proteins” option to analyze the preconstructed protein data without the ab initio gene-modeling step.

Constructed gene models were annotated by several in silico searches. Gene functions were predicted based on BLASTp version 2.2.31 (Database = TrEMBL, Swiss-Prot, and “S288C_reference” on the SGD database) and summarized by AHRD version 3.3.3 (https://github.com/groupschoof/AHRD). Protein motifs were searched using a Pfam analysis with InterProScan ver. 5.36-75 [[11]](https://paperpile.com/c/bDgfmN/6y4h). The functional ID (KEGG ID) of the proteins was determined using KofamKOALA [[12]](https://paperpile.com/c/bDgfmN/3nNy). Orthologous relationships were classified by SonicParanoid version 1.0 using the data of other fungal species (Supplementary Data 2).

Contamination identification: We eliminated the sequences derived from contaminated DNA during the sample preparation based on the Illumina read depth of coverage and homology search. We calculated the average read depth of coverage from the bowtie2 mapping of Illumina data (DRR121633 and DRR121632) against the assemblies and considered assemblies showing lower than 10× coverage as possible contaminants. We then predicted the origin of the genes on the assemblies using a BLASTp search against the refseq_genomic database and MEGAN5 [[13]](https://paperpile.com/c/bDgfmN/NL2T) and excluded the assemblies that encoded fungal genes from the possible contaminants. Our contamination identification detected 108 assemblies as contaminants (Supplementary Data 8). After the elimination of the contigs, we isolated a complete mitochondrial sequence from the contigs and then submitted the assemblies to the DDBJ as whole-genome shotgun sequence data of *R. clarus* HR1 (nuclear DNA = BLAL01, mitochondrial DNA = LC506577). For the prompt submission of the genomic gene model data to DDBJ, we used GFF2MSS ver. 3.0.2 script (<https://github.com/maedat/GFF2MSS>).

<Total RNA isolation methods based on Nakagawa et al. (2011) [[14]](https://paperpile.com/c/bDgfmN/3RsM)>

Harvested thalli (approximately 200 mg) were ground to a fine powder in liquid nitrogen, immediately dissolved in 800 µl of extraction buffer, and incubated at 65°C for 10 min. The extracts were passed through QIA shredder (Qiagen) to fractionate genomic DNAs. After two successive extractions with chloroform, RNAs were precipitated using 2.5 M LiCl at –20°C overnight. The precipitated RNAs were collected by centrifugation, dissolved in 100 µl of water, and further purified by ethanol precipitation.

<“Full-Length 16S Amplification, SMRTbell Library Preparation>

We used the PrimeScript 1st strand cDNA Synthesis Kit (TAKARA, Japan) for reverse transcription, KAPA HiFi HotStart ReadyMix (KAPA Biosystems) for DNA amplification, and SMRTbell Template Prep Kit 1.0 (Pacific Biosciences) for adapter ligation. A custom primer set was used for the reverse transcription and the first-round DNA amplification. Barcoded Universal F/R Primers Plate-96 (Pacific Biosciences) was then adopted for the second-round amplification.

Primer data

| ID | Sequence |
| --- | --- |
| AM_riboAmp_F2 | /5AmMC6/cagtcgaacatgtagctgactcaggtcacCGCTGAACTTAAGCATATCAATAAGC |
| AM_riboAmp_R1 | /5AmMC6/tggatcacttgtgcaagcatcacatcgtagCTATTGCAACAACACTTCATCAGTAG |

Lowercase = Universal sequence for the library construction

Uppercase = Designed sequence for *R. irregularis* rRNA

1st step PCR

[95ºC, 5 min.]

↓

[95ºC, 30 sec.→64ºC, 30 sec.→72ºC, 10 min.] x 10 cycles

2nd step PCR

[95ºC, 5 min.]

↓

[95ºC, 30 sec.→57ºC, 30 sec.→72ºC, 10 min.] x 20 cycles

Reference

1.	Maeda T, Kobayashi Y, Kameoka H, Okuma N, Takeda N, Yamaguchi K, et al. Evidence of non-tandemly repeated rDNAs and their intragenomic heterogeneity in Rhizophagus irregularis. *Commun Biol* 2018; **1**: 87.

2.	Koren S, Walenz BP, Berlin K, Miller JR, Bergman NH, Phillippy AM. Canu: scalable and accurate long-read assembly via adaptive k-mer weighting and repeat separation. *Genome Res* 2017; **27**: 722–736.

3.	Chin CS, Alexander DH, Marks P, Klammer AA, Drake J, Heiner C, et al. Nonhybrid, finished microbial genome assemblies from long-read SMRT sequencing data. *Nat Methods* 2013; **10**: 563–569.

4.	Walker BJ, Abeel T, Shea T, Priest M, Abouelliel A, Sakthikumar S, et al. Pilon: an integrated tool for comprehensive microbial variant detection and genome assembly improvement. *PLoS One* 2014; **9**: e112963.

5.	Langmead B, Salzberg SL. Fast gapped-read alignment with Bowtie 2. *Nat Methods* 2012; **9**: 357–359.

6.	Hoff KJ, Lomsadze A, Borodovsky M, Stanke M. Whole-Genome Annotation with BRAKER. *Methods Mol Biol* 2019; **1962**: 65–95.

7.	Kim D, Paggi JM, Park C, Bennett C, Salzberg SL. Graph-based genome alignment and genotyping with HISAT2 and HISAT-genotype. *Nat Biotechnol* 2019; **37**: 907–915.

8.	Slater GS, Birney E. Automated generation of heuristics for biological sequence comparison. *BMC Bioinformatics* 2005; **6**.

9.	Hoff KJ, Stanke M. WebAUGUSTUS—a web service for training AUGUSTUS and predicting genes in eukaryotes. *Nucleic Acids Res* 2013; **41**: W123–W128.

10.	Simao FA, Waterhouse RM, Ioannidis P, Kriventseva EV, Zdobnov EM. BUSCO: assessing genome assembly and annotation completeness with single-copy orthologs. *Bioinformatics* 2015; **31**: 3210–3212.

11.	Jones P, Binns D, Chang H-Y, Fraser M, Li W, McAnulla C, et al. InterProScan 5: genome-scale protein function classification. *Bioinformatics* 2014; **30**: 1236–1240.

12.	Aramaki T, Blanc-Mathieu R, Endo H, Ohkubo K, Kanehisa M, Goto S, et al. KofamKOALA: KEGG ortholog assignment based on profile HMM and adaptive score threshold. *Bioinformatics* 2019.

13.	Huson DH, Beier S, Flade I, Górska A, El-Hadidi M, Mitra S, et al. MEGAN Community Edition - Interactive Exploration and Analysis of Large-Scale Microbiome Sequencing Data. *PLoS Comput Biol* 2016; **12**: e1004957.

14.	Nakagawa T, Kaku H, Shimoda Y, Sugiyama A, Shimamura M, Takanashi K, et al. From defense to symbiosis: limited alterations in the kinase domain of LysM receptor-like kinases are crucial for evolution of legume-Rhizobium symbiosis. *Plant J* 2011; **65**: 169–180.
