## Supplementary Data 9 for "Conservation and host-specific expression of non-tandemly repeated heterogenous ribosome RNA gene in arbuscular mycorrhizal fungi"

### **Supplementary Data** **Legends**

**Supplementary Figure 1.**  Expanded view of Fig. 1a. Orange colors mean distance between the contig edge and the 48S rDNA region.

**Supplementary Figure 2.**  Predicted rDNA copy number and the distribution on the constructed genome data of the 18S and 28S rDNA. This figure includes the omitted species (seven non-AM fungal species; Jfl, <i>*Jimgerdemannia flammicorona*</i>; Lco, <i>*Lichtheimia corymbifera*</i>; Sra, <i>*Syncephalastrum racemosum*</i>; Mel, <i>*Mortierella elongata*</i>; Ltr, <i>*Lobosporangium transversale*</i>; Lpe, <i>*Linderina pennispora*</i>; Cre, <i>*Coemansia reversa*</i>) in Fig 2. In these seven species, multiple rDNA genes were not detected in the public genome data. The tree on the left was generated by the ML analysis of 96 single-copy genes.

**Supplementary Table 1.** Sequence data used for the *R. clarus* genome assembly.

**Supplementary Table 2.** Assembly statistics of *R. irregularis* and *R. clarus* genomes.

**Supplementary Data 1.** Start and end positions of the determined rDNA regions.

**Supplementary Data 2.** Adopted SRA data set for rDNA number count and ortholog analysis.

**Supplementary Data 3.** The number of variations between the rDNA paralogs.

**Supplementary Data 4.** Data set for PacBio Circular Consensus Sequencing (CCS) and obtained read statistics.

**Supplementary Data 5.** DRA (RNA-seq) data used for the gene model construction of *R. clarus*.

**Supplementary Data 6.** Constructed gene models from our *R. clarus* genome and their annotation information.

**Supplementary Data 7.** Comparison of the constructed *R. clarus* gene models with previously released genomic data.

**Supplementary Data 8.** Contamination identification in the constructed contigs form *R. clarus* DNA.

**Supplementary Data 9.** Primer and PCR reaction data for PacBio CSS library construction.
