## Supplementary Figure 1 for "Conservation and host-specific expression of non-tandemly repeated heterogenous ribosome RNA gene in arbuscular mycorrhizal fungi"

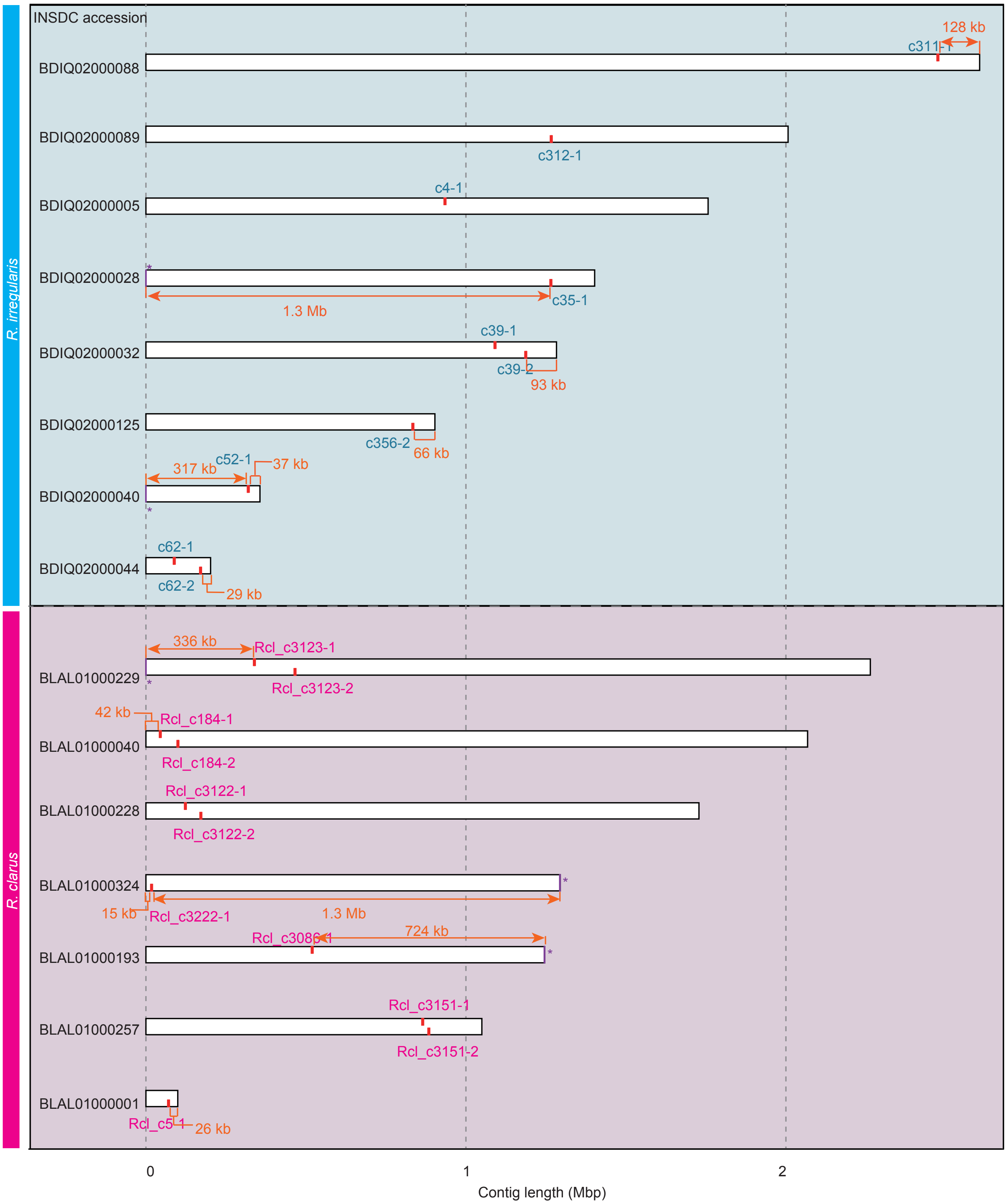


**Supplementary Figure 1.**

Expanded view of Fig. 1a. Orange colors mean distance between the contig edge and the 48S rDNA region.
