## Supplementary Figure 2 for "Conservation and host-specific expression of non-tandemly repeated heterogenous ribosome RNA gene in arbuscular mycorrhizal fungi"

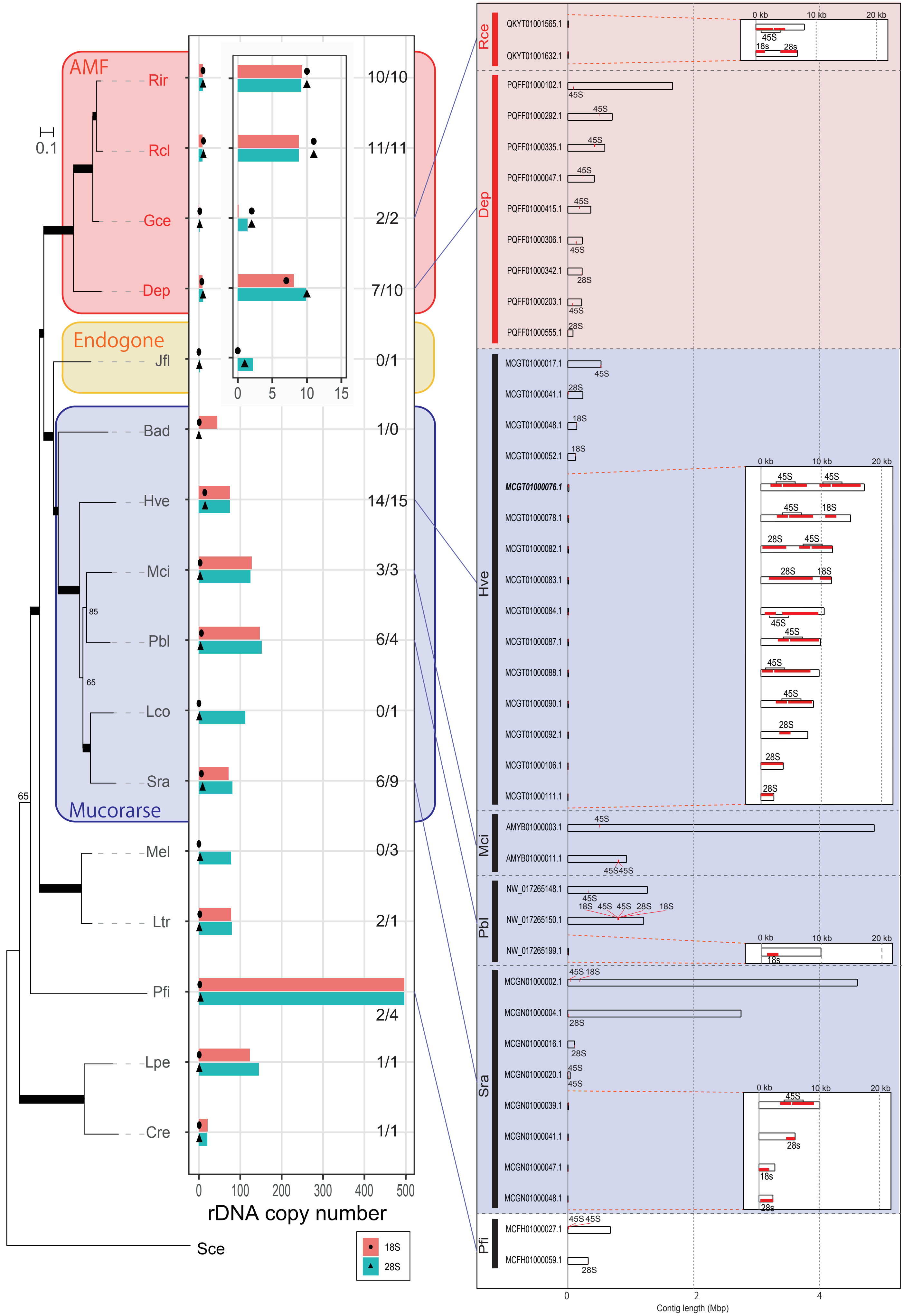


**Supplementary Figure 2.** Predicted rDNA copy number and the distribution on the constructed genome data of the 18S and 28S rDNA. This figure includes the omitted species (seven non-AM fungal species; Jfl, *Jimgerdemannia flammicorona*; Lco, *Lichtheimia corymbifera*; Sra, *Syncephalastrum racemosum*; Mel, *Mortierella elongata*; Ltr, *Lobosporangium transversale*; Lpe, *Linderina pennispora*; Cre, *Coemansia reversa*) in Fig 2. In these seven species, multiple rDNA genes were not detected in the public genome data. The tree on the left was generated by the ML analysis of 96 single-copy genes.
